# Transcranial Magnetic Stimulation in Awake Rhesus macaques: Validation of a Novel Non-invasive Apparatus

**DOI:** 10.64898/2026.03.31.715530

**Authors:** Anna Padányi, Balázs Knakker, Evelin Kiefer, Balázs Lendvai, István Hernádi

## Abstract

Transcranial magnetic stimulation (TMS) is a non-invasive brain stimulation technique widely employed in basic and clinical research. Non-human primates (NHPs) represent translationally valuable models due to their close anatomical and functional similarity to humans. However, significant technical challenges remain in implementing human-like TMS protocols in awake NHPs. Here we developed a non-invasive head- and arm-fixation apparatus that enables reliable stimulation and electromyography recordings in awake NHPs without surgical intervention and validated the apparatus with two TMS protocols in rhesus macaques. First, we implemented an adaptive motor threshold (MT) determination method developed recently for humans, which converged successfully to valid MTs as defined by the International Federation of Clinical Neurophysiology. Second, we measured a robust short-interval intracortical inhibition effect for the first time in awake NHPs. Successful implementation of human TMS protocols in awake NHPs provides proof-of-concept validation of our apparatus, paving the way to bidirectionally translatable, clinically relevant neuromodulation protocols.

## 1. Introduction

Transcranial magnetic stimulation (TMS) is a non-invasive brain stimulation (NIBS) technique that has become a valuable tool in both basic neuroscience research and clinical practice. By inducing brief magnetic pulses non-invasively over the scalp, TMS enables us to probe and modulate cortical excitability. Despite the substantial technical progress and numerous novel protocols in the last forty years of TMS, our knowledge on the underlying mechanisms of the neuromodulatory effects of TMS remain limited. While rodent models offer mechanistic insights into brain circuitry, their anatomical and physiological dissimilarity from humans poses constraints on generalisability of the data, highlighting the need to bridge the translational gap between preclinical animal experiments and clinical (human) studies. Non-human primates (NHPs), particularly macaques, are ideal models for bridging this translational divide^1^, given the high degree of structural and functional homology with the human brain, including direct cortico-motoneuronal projections to the hand and the similar anatomical organisation, comparable thickness^2^ and gyrification^3^ of the neocortex with similar functional output.

However, the application of TMS in NHPs presents technical challenges. While conducting experiments under anaesthesia prevents movement of the animals and enables consistent and accurate stimulation, anaesthesia influences the cortical state itself. In awake state, surgically implanted headposts are widely used, which similarly prevent head movement and allow precise targeting, hence accurate and consistent TMS application. However, as surgical implantation of a headpost is not without its own challenges, head fixation methods without surgical intervention offer a good, alternative, non-invasive solution enabling TMS in conscious NHPs in a reliable and replicable manner. Additionally, non-invasive protocols allow direct translation from humans to NHPs and backwards. Nonetheless, only a small number of groups have develop non-invasive head fixation methods for TMS to date^1,4–7^, and their implementation is often highly dependent on the exact set-up of the given laboratory.

In response to these challenges, here we introduce a novel, non-invasive head- and arm-fixation apparatus design to accommodate different cranial morphologies and minimise movement during stimulation. The head-fixation unit also allows for flexible TMS coil positioning, enabling access to a range of cortical regions while preserving the animal’s comfort and alertness. Additionally, the apparatus incorporates an arm fixation unit, restricting movement of the fingers, the lower and the upper arm, allowing for electromyographic (EMG) recording from the muscles of the hand, particularly the abductor pollicis brevis (APB) muscle.

As a proof-of-concept, we first measured the motor threshold (MT) according to an adaptive stepping protocol^8^ in 4 awake rhesus macaques. Second, for the first time in awake NHPs, a paired-pulse protocol (short-interval cortical inhibition, SICI) was applied in 2 subjects. By this, we address the technical challenges of neuronavigation-assisted TMS application in awake NHPs with a novel, adjustable apparatus and provide the basis of highly translatable TMS research on awake, surgically intact non-human primates.

## 2. Methods

### 2.1. Animals and housing

This study was performed using 4 adult male rhesus macaque monkeys (*Macaca mulatta*), aged 14.2 ± 3.3 years (ranging from 10.7 to 18.6 years), weighing 10.1 ± 0.6 kg (ranging from 9.2 to 10.6 kg). The macaques were chair-trained subjects, who had been previously accustomed to a cognitive behavioural set-up and participated in various cognitive paradigms.

As also described in our previous studies^9^, subjects were housed in pairs in home cages that were uniformly sized 230 × 100 × 200 cm (height × width × depth) and were equipped with wooden rest areas. In the vivarium and laboratories, temperature and humidity were maintained at 24 ± 1°C and 55 ± 5 RH% with continuous airflow. Light conditions were set to a 12/12 h light-dark cycle (lights on at 07:00) using full spectrum lighting with transitions (twilight periods) between light and dark phases and were supplemented by natural illumination through side and roof windows to maintain the natural rhythm of the subjects. Subjects were fed once per day between 3 and 4 PM, following the daily training or testing sessions. Diet was standard nutritionally complete laboratory chow specially designed for non-human primates (Altromin Spezialfutter GmbH) and was daily supplemented with fresh fruit and vegetables.

All procedures were conducted in the Grastyán Translational Research Centre. This study was approved by the Animal Welfare Committee of the University of Pécs and the Hungarian National Scientific Ethical Committee on Animal Experimentation, and the ethical permission was issued by the Department of Animal Health and Food Control of the County Government Offices of the Ministry of Agriculture (BA02/2000-25/2020). Measures were taken to minimize pain and discomfort of the animals in accordance with the Directive 40/2013. (II.14.): ‘On animal experiments’ issued by the Government of Hungary, and the Directive 2010/63/EU ‘On the protection of animals used for scientific purposes’ issued by the European Parliament and the European Council.

### 2.2. Novel apparatus

#### 2.2.1. Non-invasive head-fixation unit for TMS

A 3-piece TMS head-fixation apparatus was developed to minimise head movement while leaving the frontal, parietal and part of the occipital regions available for TMS stimulation. A 3D-printed, static base (**Figure 1B, a**), prepared with Fused Deposition Modelling (FDM) from Polyethylene Terephthalate Glycol-Modified (PETG), can be mounted to the neck plate of a generic primate chair (**Figure 1A**) and adjusted with four screws hidden in grooves so that the base may be moved parallel to the neck plate to ensure comfortable posture for the subjects. The back piece (**Figure 1B, b**) and the front piece of the head-fixation apparatus, the latter formed by the mask piece (**Figure 1B, c**) and the handle (**Figure 1B, d**), are attachable to the base. The back piece and the handle were 3D-printed with FDM technology from Thermoplastic Polyurethane (TPU). The back piece can be adjusted by shifting in the horizontal plane and fixed by four hand screws (**Figure 1B, e**), 2 on each side. The back piece has a head cushion (**Figure 1B, f**). The front piece has an inner layer, a face mold (**Figure 1B, g**) made of two-component, high tensile strength silicon (Shore 30A hardness). Utilising the 3D face reconstructions of the subjects based on MRI images of the subjects (see below), face molds of three different sizes (small, medium and large) were fabricated with appropriately sized opening and angles to follow the cheekbones as closely as reasonable.

**Figure 1:**
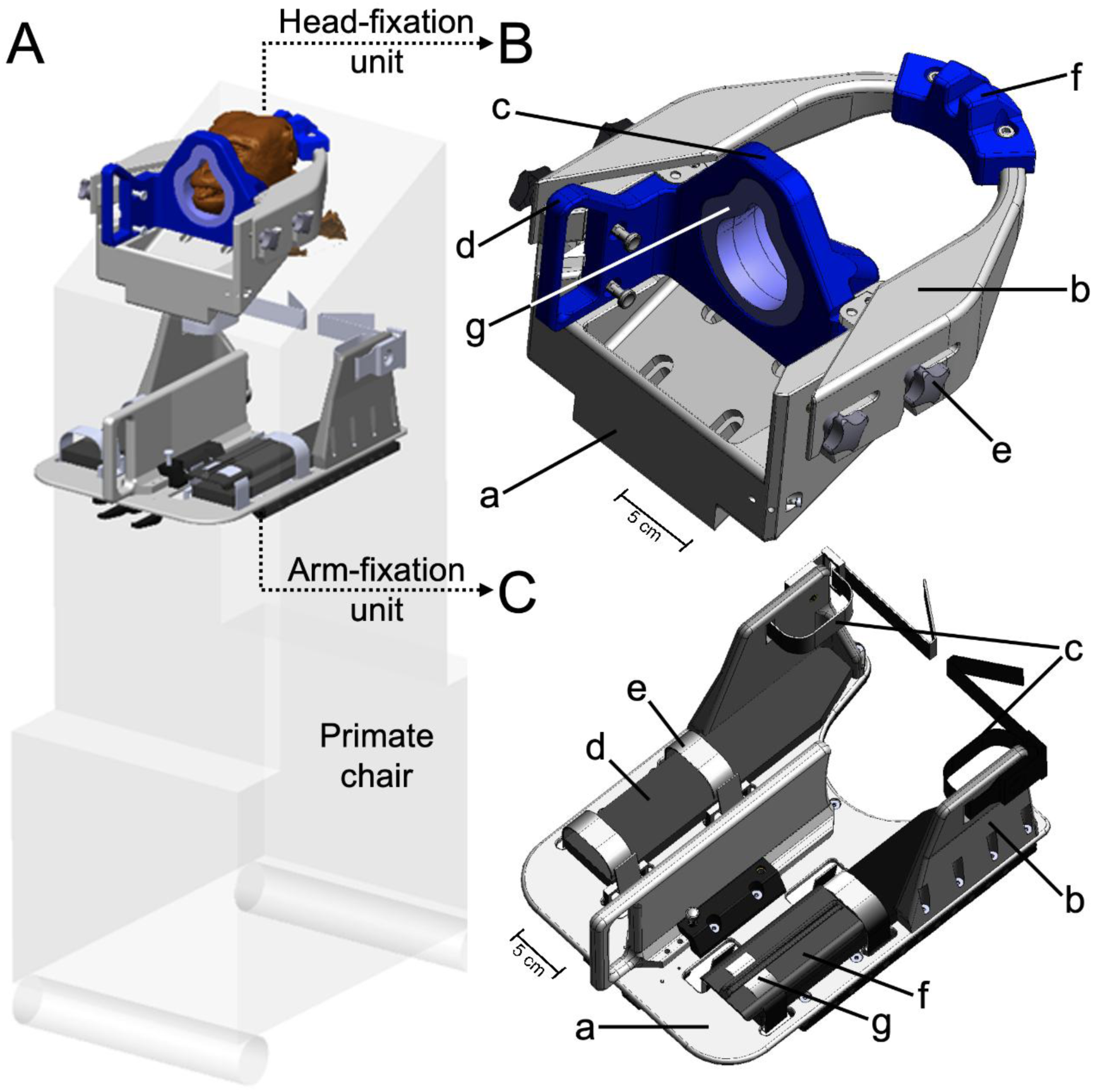
The novel apparatus developed for facilitating frameless neuronavigation assisted transcranial magnetic stimulation (TMS) and electromyography (EMG) in awake rhesus macaques. **A:** The apparatus designed to be attached to a primate chair comprises two primary components: the head-fixation unit and the arm-fixation unit. **B:** The head-fixation unit – minimising head movement during stimulation – consists of three main parts: the static base (a), the detachable back piece (b) and the detachable front piece consisting of the mask (c) and the handle (d). The hand screws (e), head cushion (f) and face mold (g) make the head-fixation unit adjustable and customised for improved subject comfort. **C:** The arm-fixation unit – holding either one arm or both arms, including the fingers – enables reliable EMG recordings from hand muscles. It consists of a horizontal plane (a) onto which the ankle holders (b) and the foam platform (d) are mounted. Fastening bands are holding the upper (c) and lower arm (e) in place with the fingers further held by a plate (f) and its fastening (g). (Blueprint renderings courtesy of ADOTT Solutions Ltd.) [figure width: 17.3 cm]

#### 2.2.2. Arm-fixation unit for EMG recording

Additionally, an arm-fixation apparatus, i.e. arm holder, was developed to restrain the arm and the fingers separately – leaving muscles of the hand accessible for electromyography (EMG) recording to enable the measurement of motor-evoked potentials (MEPs) in response to single-pulse TMS. The arm holder is attached to a horizontal plate made of an aluminium alloy (EN AW-5083, **Figure 1C, a**) with 3D-printed ankle holders (**Figure 1C, b**) prepared with FDM from PETG. Polyamide fastening bands (**Figure 1C, c**) run through the side to restrict movements of the upper arm, keeping the arm at the ankle holder. The lower arm rests on a 15-mm thick foam platform (**Figure 1C, d**) made of ethylene-propylene-diene monomer (EPDM), and the movements of the lower arm are also restricted by a polyamide fastening ending in Velcro (**Figure 1C, e**). The fingers are held by a 3D-printed plate (**Figure 1C, f**) fixed once more by polyamide Velcro fastening (**Figure 1C, g**).

### 2.3. Validation of the novel apparatus by TMS-MEP

The preparatory phase of each session included the application of the TMS head- and arm-fixation apparatus as described above. The right palm of the subject was wiped with alcoholic disinfectant before disposable surface EMG electrodes (30 mm diameter, Ag/AgCl) were placed above the abductor pollicis brevis (APB) muscle of the right thumb in a tendon-belly montage to record MEPs. The ground was attached to the primate chair. Raw EMG waveform data was recorded in AC mode at 3 kHz sampling rate, digitised at 12 bit with an analogue bandpass filter set between 16–470 Hz (EMG Pod, Rogue Research Inc., Montreal, Canada). Data was saved in epochs between −50ms to +150ms relative to TMS pulse onset time (Brainsight™ TMS Frameless Navigation system, Rogue Research Inc, Montreal, Canada). Peak-to-peak amplitude of the MEPs in the 10 ms to 90 ms time window following stimulation was calculated by the Brainsight software.

To induce the MEPs, focal biphasic magnetic stimuli were applied over the scalp using a MagPro x100 including MagOption (MagVenture, Lucernemarken, Denmark) via a figure-eight coil with active cooling (Cool-B65, MagVenture). The coil was manually held and placed tangentially over the hand area of the left motor cortex at an angle of 45 degrees to the sagittal plane, inducing a postero-anterior directional current flow.

Prior to the start of the experiments each subject underwent a T1-weighted MRI (Siemens Magnetom Prisma Fit and 16-channel Pediatric head coil, Siemens Healthineers AG, Erlangen, Germany; 3T, 3D MPRAGE with isotropic 0.6 mm spatial resolution; TR/TE/FA = 2700 ms/2.19 ms/9°; FOV = 153.6 × 192 × 192 mm^3^) scan in order to create the individual stereotaxic map of their brain. Brain images were imported into the Brainsight™ TMS Frameless Navigation system (Rogue Research Inc, Montreal, Canada) wherein head models were created to allow for continuous online control of coil positioning during the experimental sessions using optical tracking of fiducial markers on the coil and on the head-fixation apparatus of the subject.

The APB muscle area was identified in the primary motor cortex (M1 hotspot) for each subject by systematic biphasic single-pulse stimulation pattern covering an area centred around the expected M1 location on the precentral gyrus within 8-10 mm in parallel to the precentral gyrus and 15-20 mm perpendicular to the precentral gyrus. The location yielding the largest and most consistent MEPs at a stable stimulation intensity (SI) was marked in the neuronavigation system and was further referred to as ‘M1 hotspot’. At the beginning of all sessions, the same M1 hotspot location was targeted, verified with a short, asterisk-shaped stimulation pattern in the 4 cardinal directions and the 4 intermediate directions relative to the coil orientation.

Data analysis and visualisation was conducted offline using R 4.3.2 (R Core Team, 2024) in RStudio (Posit Team, 2025).

#### 2.3.1. Adaptive motor threshold measurements

Traditionally, the motor threshold (MT) is defined as the minimum stimulation intensity at which the MEP peak-to-peak amplitude exceeds a criterion level (usually set between 50–100 µV) in 50% of the trials^10,11^. This MT is often measured with the relative-frequency method^8,12^. However, that usually requires a substantial amount of pulses delivered, while newer methods, like the adaptive-stepping paradigm^8^ can determine MT in 25 pulses with high confidence. Minimizing the number of pulses has benefits not only in clinical settings, but also in translational research with rhesus macaques from both animal welfare and experimental design perspective.

We measured the MT based on the 100 µV criterion^6,9,10^ in all 4 subjects (see *Animals and housing*) using DCS-HA (Digital Control Sequence – Harmonic, Adaptive), a stochastic approximation method employing a stimulation sequence with adaptive stepping which follows harmonic convergence^8^. The initial step size set in the algorithm is 4.2% of Maximum Stimulator Output (%MSO), which decreased to 1 %MSO after 6-9 steps. In the adaptive control sequence, step size is only decremented if the MEP amplitude changes from suprathreshold to subthreshold, or vice versa. Specifically, we used the DCS-HA algorithm implemented in the browser-based Stochastic Approximator of MT (SAMT) software by Wang and colleagues^8^ (https://tms-samt.github.io/, Version 1.7.3 or 1.7.4 – the versions only differed in user interface, the algorithm was identical), which after manual entry of stimulation intensity and its result (suprathreshold response present or absent), outputted the current threshold estimate and next stimulation intensity that we used (also adjusting manually), repeating until the software indicated convergence (at pulse number 25 in the versions we used).

Based on simulations^8^ and human clinical data^13^ , the algorithm produces a reliable MT estimate after 20-25 pulses, given that the number of suprathreshold responses for the last 10 pulses (n-of-ten) is between 3 and 7 (inclusive). Besides this built-in convergence criterion in SAMT, to further validate the threshold estimate, we visualised the empirical distribution of peak-to-peak amplitudes to perithreshold stimulation intensities in the convergence window. The integer %MSO levels below and above the estimated MT were selected as peri-threshold intensities, because the MT estimate was provided to 1 decimal figure precision by the app, but the stimulator could only be adjusted with 1 %MSO steps. Since the algorithm uses the n-of-ten metric to evaluate convergence which is expected to occur the earliest at the 20^th^ pulse^13^, the convergence window was defined to start from pulse number 10.

It should be noted that after we had acquired the data, newer versions of the SAMT software^13^ have been published that already start checking for convergence at pulse number 20, not 25 as in our version. Based on our data, we know that this would have meant that during our MT measurements the newer versions of the software would have indicated convergence already at pulse No. 20. However, we are the first to explore the applicability of this method in rhesus macaques (as a proof-of-concept validation of our novel apparatus), the behaviour of the algorithm and how the expected accuracy of the MT estimate decreases with more pulses is not yet known, so as a conservative strategy, we kept using the estimate at pulse No. 25.

#### 2.3.2. Short-interval cortical inhibition (SICI)

SICI was assessed using a paired-pulse TMS protocol delivered over the primary motor cortex (M1) of the left hemisphere of 2 subjects (‘Sp’ and ‘Sz’). Stimuli were delivered with the same experimental set-up as for adaptive MT described above (see *Validation of novel apparatus by transcranial magnetic stimulation*). Each conditioned, or paired-pulse (pp) trial consisted of a subthreshold conditioning stimulus (CS) followed by a suprathreshold test stimulus (TS), with interstimulus intervals (ISI) of 1, 2.5 and 5 ms. The intensity of the CS was set to 80% of the subject’s MT (80 %MT), while the TS was set to 120 %MT. Unconditioned single-pulse (sp) trials consisted of a single stimulation pulse at the same level as the TS (120 %MT). From both subjects, 2 sessions were recorded with 4-5 stimulation blocks in each. One stimulation block consisted of 9 paired-pulse (pp, 3 from each ISI) and 9 single-pulse (sp, unconditioned) trials delivered in an alternating sp-pp sequence. The blocked design was used to account for potential fluctuations of overall MEP amplitudes. Paired-pulse ISI was pseudorandomised in each block and intertrial interval was kept at the median of 7 s with ± 1 s jitter. MEP segments with major artefacts or subject misbehaviour were manually marked and excluded from analysis.

SICI was quantified as the ratio (expressed in percentages) of MEP amplitude in the paired-pulse condition relative to the mean unconditioned MEP amplitude within each block: SICI = conditioned V_p-p_ / unconditioned V_p-p_, where V_p-p_ is peak-to-peak MEP amplitude. Peak-to-peak amplitudes were log-transformed, and all subsequent descriptive and inferential statistics were calculated on these transformed logV_p-p_ values. This is considered best practice due to the lognormal distribution of MEP amplitudes and its multiplicative scaling^9,14^. We applied linear models to the transformed values, wherein the contrasts formed as conditioned minus unconditioned logV_p-p_ estimate the log-transformed version of the above-defined SICI ratio, and statistical tests on the log scale against 0 probe whether the ratio differs from 100%. In text description and visualisation (all expected values and confidence intervals shown in Figure 3 or described in the SICI results text), statistical results are back-transformed to the original linear ratio scale: arithmetic means and marginal means from linear models on log-scale data correspond to geometric means on the linear ratio scale, and confidence intervals estimated on the log scale represent multiplicative uncertainty on the linear ratio scale.

**Figure 2:**
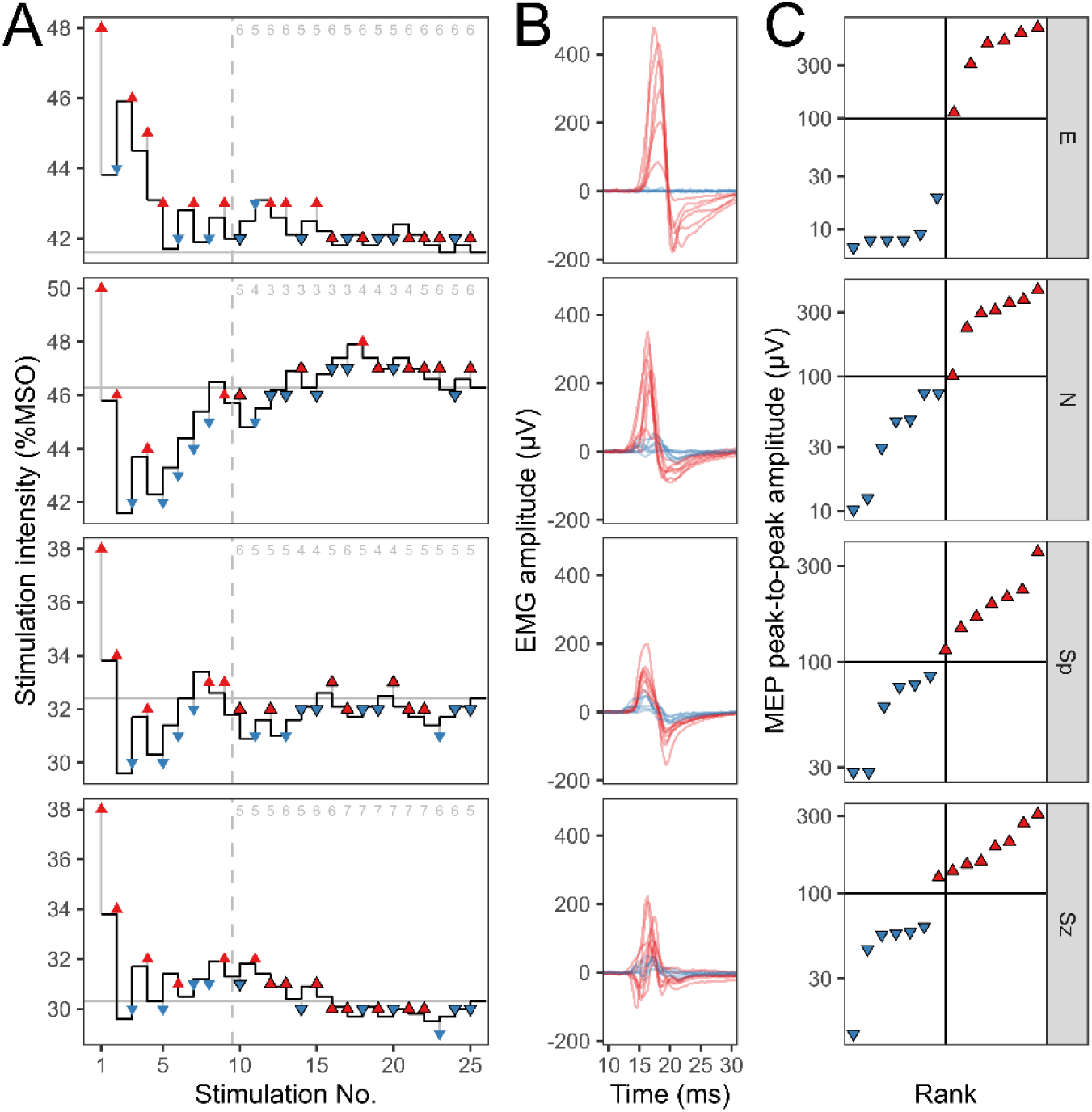
Single sessions of resting motor threshold measurement with our novel, adjustable, non-invasive apparatus in 4 awake rhesus macaques. **A)** Stimulation intensity sequence (y coordinate of the triangle markers) determined on-line using the DCS-HA method ^8^. Updates of the MT estimate after each pulse are shown by the steps of the black line, with subthreshold responses (blue downward triangles) increasing, and suprathreshold responses (red upward triangles) increasing the estimate. The determined final MT is shown by a horizontal gray line). The dashed vertical line shows the start of the convergence window (see Methods). For perithreshold pulses (integer %MSO below and above MT in the convergence window) the triangle markers have a black outline, these are included in **B** and **C**. Gray numbers at the top indicate the number of suprathreshold responses in a sliding window of the last 10 TMS pulses. **B)** MEP signals arising from perithreshold pulses displayed with colour indicative of the peak-to-peak amplitude in relation to the 100µV threshold (blue below, red above). **C)** Empirical distribution of response amplitudes to perithreshold pulses demonstrating 50% suprathreshold response rate, as expected by the definition of the MT. The pulses are sorted along the x axis according to peak-to-peak amplitude (y axes), the threshold value of 100 µV is shown as a horizontal black line, and the 50^th^ percentile of the distribution is shown as a vertical black line. Suprathreshold responses are shown with red markers, subthreshold responses with blue markers. [figure width: 11.8 cm]

**Figure 3:**
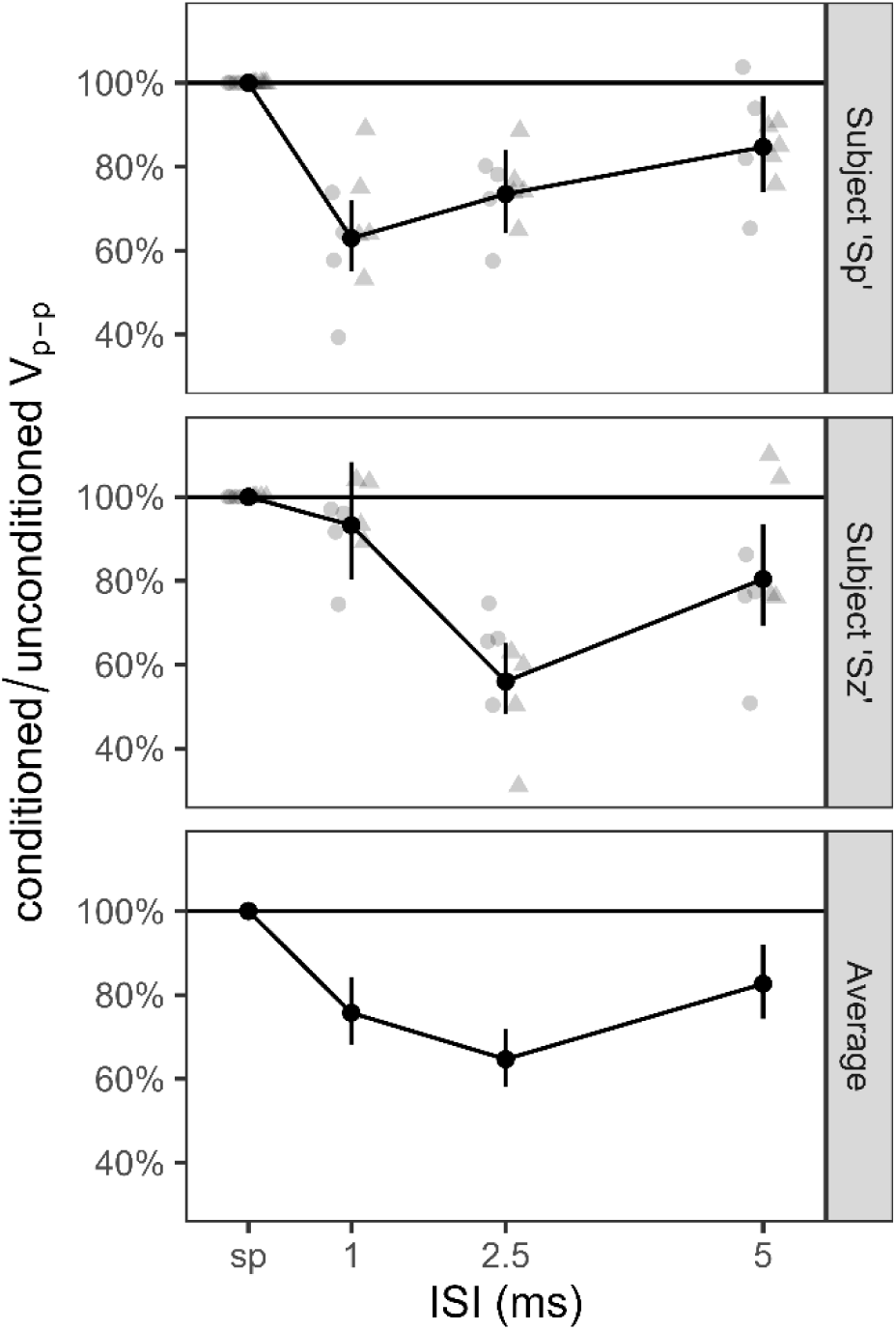
Ratio of peak-to-peak MEP amplitudes (y axes) for conditioned over unconditioned stimulations for paired pulse stimulations with 1, 2.5 and 5 ms interstimulus intervals (x axes) in 2 sessions in 2 rhesus macaques analysed separately in a per-subject model (top panels, labelled subject ‘Sp’, Subject ‘Sz’), and in an aggregate model (bottom panel, ‘Average’). Light grey markers on per-subject panels indicate blockwise (geometric) mean response amplitude ratios of unconditioned, control single pulses and conditioned, i.e. paired pulses; shapes of the markers differentiate the 2 sessions. The horizontal black line indicates 100%, which is the average of MEP amplitude level of the unconditioned (single) pulses. Black shapes indicate expected values of MEP amplitude ratios for each ISI with whiskers denoting 95% simultaneous confidence intervals based on the respective per-subject or aggregate models. Note that though only block-level summaries are displayed on the figure, the means and confidence intervals are (hierarchical estimates) based on individual pulse response amplitudes. [figure width: 8.2 cm]

To provide an estimate of the SICI effects from all the data (altogether 4 sessions from 2 subjects) logV_p-p_ was modelled using a linear mixed model (LMM, in lme4 R package version 1.1.37) with a fixed effect for ‘Stimulation condition’ as a dummy-coded categorical variable (sp – intercept, pp 1 ms, pp 2.5 ms, pp 5 ms), and nested random intercepts for blocks, sessions and subjects. The primary outcomes of interest were the SICI effects, that is, each pp ISI condition compared to the sp control, as also defined above. For the contrasts estimated for these effects in the LMM, the single-step multivariate-t-based method (as implemented in the emmeans R package as procedure “mvt”, but see also Bretz and colleagues^15^) was used for multiplicity correction, yielding both p-values (denoted p_c1_) and simultaneous confidence intervals. After establishing whether the effect is present, a separate family of tests with the same correction method was conducted for pairwise comparisons of pp ISI conditions to probe how the SICI effect depends on ISI (with the resulting p-values denoted as p_c2_). Degrees of freedom was approximated using the Kenward-Roger method in all tests (using R packages lmerTest and emmeans).

Single-subject LMMs analogous to the main model were also fit with random intercepts for blocks nested in sessions. For these single-subject LMMs, we describe and visualise marginal means and simultaneous confidence intervals using the single-step method in the emmeans R package^15^. Expected values and confidence intervals on the single-subject panels of **Figure 3** are from these models. In addition, for the specific purpose of assessing the similarity between the SICI effects of the 2 sessions, we fitted single-subject linear models (termed session consistency models) on the block geometric means with a categorical factor for session, the Stimulation condition factor, and their interaction. The interaction in this model quantifies how consistent the SICI pattern is between the 2 sessions in the 2 subjects.

## 3. Results

We have developed a novel, adjustable, non-invasive head- and arm-fixation apparatus for TMS stimulation and MEP recording from the hand in awake macaques. To validate our apparatus, we present data from two widely used TMS paradigms: the obligatory initial protocol, motor threshold (MT) measurement and short-interval cortical inhibition (SICI), a paired-pulse protocol.

### 3.1. Habituation to the novel apparatus

Subjects were habituated to the apparatus by positive reinforcement training. A single habituation training session was conducted each day, usually taking place between 8:00 and 13:00, with individual sessions lasting between 10 to 40 min. Training proceeded in stages. First, subjects were accustomed to the back and the front pieces of the head-fixation apparatus (**Figure 1**) separately, then together. On average, this first habituation step took 11±7.6 (mean±SD) sessions (ranging from 5 to 24 sessions). Then, subjects were habituated to the arm fixation apparatus, progressively increasing the number of the fastening bands. This step was only necessary for 2 out of 4 subjects and lasted for 2.5±1.5 sessions on average. Finally, the full set-up was applied to the subjects, and during an additional habituation phase of 4.8±1.5 sessions (ranging from 3 to 7 sessions), the time spent sitting calmly in the set-up was gradually increased up to approximately 20 min. The full training took 16.0±8.8 (ranging from 9 to 31) sessions for all 4 subjects.

### 3.2. Motor threshold determination using an adaptive stepping method

Using our novel head- and arm-fixation apparatus, we first measured resting MT (as defined by the International Federation of Clinical Neurophysiology, IFCN, with 100 µV criterion^10,11^ in 4 subjects applying an online adaptive stepping algorithm implemented in the SAMT tool^8^. At pulse number 25, the MT estimate converged to 41.6, 46.3, 23.4 and 30.3 (for subjects E, N, Sp and Sz, respectively, see **Figure 1A**) with 5-6 responses in the last 10 pulses. In humans, SAMT has been shown to provide accurate MT estimates after 20-25 pulses if the number of responses per the 10 last pulses is between 3 and 7 (inclusive). In our data, this n-of-ten metric fulfilled this latter criterion between pulse No. 20 and 25. This implies that the pattern of convergence in our data is in accordance with the pattern observed in humans.

On **Figure 2B**, we show the MEP waveforms for perithreshold pulses (integer %MSO below and above MT in the convergence window), whereas **Figure 2C** shows the empirical distribution of the peak-to-peak amplitudes of these responses. Notably, while peak-to-peak amplitudes for perithreshold pulses varied in a wide range (min: 6.8 to 28.4 µV, max: 308 to 656 µV), the proportion of responses above the 100 µV threshold was around 50% (subject ‘E’: 6/12, ‘N’: 7/14, ‘Sp’: 7/13, ‘Sz’: 8/14, see **Figure 2C**), thereby fulfilling the traditional IFCN definition of MT^10,11^. Thus, we have successfully obtained valid MTs in 4 adult rhesus macaques within 25 pulses – similarly to the number of pulses required for humans, which validates our non-invasive measurement technique developed for awake non-human primates.

### 3.3. Paired-pulse stimulation – short-interval cortical inhibition

First, we analysed the aggregated data of the 2 subjects that were involved in the SICI experiment (**Figure 3**, bottom panel). Stimulation condition had a significant effect on the amplitude of MEP responses (F_3,285.0_=37.3, p<2×10^−16^): Compared to unconditioned, single pulse control stimulation (delivered at the same intensity as test stimuli: 120% MT), MEP amplitudes were smaller for conditioned stimuli with all 3 paired-pulse interstimulus intervals (ISIs). On average, the strongest effect was found for the 2.5 ms ISI: MEP amplitudes dropped to 65% of unconditioned control amplitude (95% CI: [58% 72%], t_285.0_=−9.77, p_c1_<10^−5^, corrected primary outcome) while 1 ms and 5 ms caused only smaller decrements that were still significant in the aggregated model (1 ms: 76%, [68% 84%], t_285.0_=−6.23, p_c1_<10^−5^; 5 ms: 83%, [74% 92%], t_285.0_=−6.23, p_c1_=8×10^−5^). In accordance with this, response amplitudes in the 2.5 ms condition differed from both the 1 ms (V_p-p_(2.5 ms)/V_p-p_(1 ms)=85%, t_285.0_=−2.89, p_c2_=0.011, corrected secondary outcome) and the 5 ms condition (V_p-p_(5 ms)/V_p-p_(2.5 ms)=78%, t_285.0_=−4.49, p_c2_=0.00003), but the 5 ms and 1 ms conditions did not differ from each other significantly (V_p-p_(1 ms)/V_p-p_(5 ms)=92%, t_285.0_=−1.60, p_c2_=0.25).

The top two panels of Figure 3 show data separately for the 2 subjects. Visual inspection suggests that the pattern of the SICI effects differs between the 2 subjects: subject ‘Sp’ had the strongest SICI at 1 ms, while subject ‘Sz’ showed virtually no SICI at 1 ms ISI (confidence interval on **Figure 3** overlaps with 100%) and a strong SICI at 2.5 ms ISI. The fact that the 2 sessions showed largely consistent effects within subjects (Session consistency linear model, Stimulation Condition × Session interaction, subject ‘Sp’: F_3,28_=0.56, p=0.65, subject ‘Sz’: F_3,24_=2.1, p=0.13) suggests that this indeed could reflect inter-subject but not inter-session variability.

## 4. Discussion

In the present study, we have introduced a novel non-invasive apparatus that enables the reliable and replicable application of transcranial magnetic stimulation (TMS) protocols in awake non-human primates (NHPs). Specifically, we validated our apparatus in two proof-of-concept experiments involving altogether 4 rhesus macaques: motor threshold (MT) determination using an adaptive online algorithm, and measurement of paired-pulse short-interval intracortical inhibition (SICI). Our novel apparatus thus addresses a pressing need in the field of translational neuroscience: it enables the implementation of human TMS protocols in awake NHPs under similar circumstances as in humans.

Existing approaches for NHP TMS experiments include conducting the procedure under various forms of anaesthesia^16–22^. While anaesthesia prevents voluntary movement allowing for precise TMS targeting and EMG recording without motion artefacts, the cortical state is directly influenced by the anaesthetic agents themselves. Alternatively, in awake NHPs motion is generally prevented by head fixation by a surgically implanted headpost^23–33^. While invasive head fixation also allows precise TMS targeting and the possibility of simultaneous single-cell recording in awake state^28,31^, there are still drawbacks like perioperative and long-term risks associated with anaesthesia, stressful post-operative recovery, and infection^34^. Furthermore, the protrusion of the headpost itself can impede TMS coil positioning for targeting the region of interest^34^.

Non-invasive head-fixation offers an ideal alternative solution. To date, successful non-invasive head restraint in awake macaques for TMS studies has only been reported by a handful of research groups worldwide. Yamanaka and colleagues^7^ employed a mesh-like thermoplastic mask custom-fitted for each subject, and Honda and colleagues^5^ developed an alternative thermoplastic mask with urethane cushion for comfort. Testing some of these thermoplasts, we have faced difficulties due to the limited geographical availability of the vendor, the challenging process of mask molding, and the inadequate head fixation of subjects with larger cranial dimensions. Therefore, we turned our focus on 3D-printable mask options, similar to that of Tang and colleagues^6^ who described a 3D-printed helmet that permits free head movement, yet in their case the helmet itself was secured invasively using in-skull titanium fixtures. As our goal was to use genuinely non-invasive technology, we developed a 3D-printable mask made of adjustable components.

In addition to minimising head movement to achieve stable and reproducible TMS delivery, the reliable recording of EMG signals from resting muscles in awake macaques is also an essential requirement for TMS-MEP experiments. For this purpose, besides our currently presented arm-fixation unit, 2 other laboratories have also developed arm-restraining systems. Amaya and colleagues^4^ used horizontally positioned Plexiglas plates with elastic bands securing the forearms and upper arms, combined with touch sensors monitoring the arm positions. Honda and colleagues^5^ proposed to loosely restrain the subject’s arm(s) in a plastic tube to enable EMG recording. While both approaches are effective, their implementation is closely linked to the set-up of the given laboratory.

In contrast, our presently introduced apparatus – including the head- and arm-fixation unit – is adjustable for accommodating different head sizes and arm lengths (for details please refer to Methods), and also for adaptation to various laboratory settings and primate chairs with minimal necessary redesigning. As our apparatus is completely non-invasive, it improves animal welfare, supports conducting longitudinal studies, and permits parallel experiments on the same animal in freely moving^35^ or social behaviour settings^36^. Furthermore, the apparatus enables stimulation in awake, unaltered cortical state, which can again facilitate pharmacological, cognitive and social behavioural experiments in combination with various TMS protocols^37^. To validate that our novel apparatus enables stimulation conditions that are directly comparable to those used in human participants, we conducted two proof-of-concept experiments.

First, we utilised the Stochastic Approximator of Motor Threshold (SAMT) tool with a DCS-HA (Digital Control Sequence – Harmonic, Adaptive) stepping protocol^8^. Simulations of human MT determination^8^ established an “n-of-ten” requirement, stating that the method yields valid MT estimates from 25 pulses when out of the 10 most recent pulses (independently of stimulation intensity), 3-7 pulses produce responses above the 100 µV peak-to-peak amplitude criterion. The number of pulses required was later lowered to 20 after testing the method on a clinical human population^13^. This n-of-ten requirement was met at our (conservative) stopping point at the 25^th^ pulse in all 4 subjects, and the requirement was even met at the newly recommended 20^th^ pulse. The data thus clearly show that in rhesus macaques SAMT yielded the same convergence pattern that had been established in humans. Besides meeting the n-of-ten requirement^8^, we also verified that our SAMT estimates in rhesus macaques fulfilled the traditional IFCN definition of MT^10,11^ using data from stimulation intensities closest to the measured threshold. It is important to note that we used the DCS-HA algorithm with the settings tailored to humans. Further optimizing the parameters of the DCS-HA algorithm for rhesus macaques – and possibly other NHP species – will require more empirical data and/or simulations. Notwithstanding, beyond the proof-of-concept validation of our novel apparatus, our results also suggest that the SAMT method can be successfully applied in rhesus macaques.

In our second experiment, using a human-relevant SICI protocol, the expected cortical inhibition effect was demonstrated for the first time in awake NHPs. The average SICI profile showed a strong inhibition effect at an interstimulus interval (ISI) of 2.5 ms and a weaker, yet detectable, inhibition effect at 1 ms. This pattern aligns with findings reported in human studies^38–41^. Interestingly, while the SICI effect was evident in both subjects, the peak of inhibition appeared reliable within, but differed strongly between subjects: subject ‘Sp’ exhibited maximal SICI at 1 ms, whereas subject ‘Sz’ showed a peak at 2.5 ms. Such variation may reflect individual differences in recruitment of interneuronal networks as suggested in humans^42^. Future studies with larger cohorts or longitudinal sampling will further characterise the reliability and variability of SICI between individual subjects. The present findings provide proof-of-concept evidence for the feasibility of SICI assessment in awake rhesus macaques, made possible by our newly designed non-invasive head- and arm-fixation apparatus.

Utilising the apparatus in a preclinical research setting also highlights possible small differences in the evoked motor responses between NHPs and humans: in Padányi et al^9^, we have previously shown that the overall shape of the recorded recruitment curves in awake rhesus macaques was in line with human data, however, the magnitude of the plateau was vastly smaller in macaques. At the same time, the SICI ratios in the present study seem to be largely consistent in range to human data^43,44^. In the present study, on average, MEP amplitudes dropped to 65% of the unconditioned control amplitude at the 2.5 ms ISI, while in humans the amplitude drop at 3 ms was between 40-60% with various inter-trial interval times^43^ or was recorded around 60%^44^.

These similarities in SICI ratio could provide us with a translational insight concerning the underlying mechanisms involving the GABAergic intracortical circuits^38–41^. Yet, further details remain a debate with arguments made for GABA_A_ receptor involvement for 2.5 – 3 ms ISI and potential influence of relative refractoriness of the inhibitory axons for 1 ms ISI^45^. Future TMS experiments using our apparatus will contribute to the comprehensive mapping of differences and similarities between NHP and human cortical excitability.

In summary, our novel head- and arm-fixation apparatus for TMS-MEP offers a robust and ethically responsible platform for conducting advanced functional neurostimulation studies in awake NHPs. Its adaptability, reliability and compatibility with human protocols mark a substantial advancement in the field of translational brain stimulation research. Its ecological validity also facilitates the reverse translation of human findings into controlled, mechanistic animal models. By enabling non-invasive and reproducible assessments of brain function, our head- and arm-fixation apparatus holds significant promise for advancing both fundamental neuroscience and the development of bidirectionally translatable, clinically relevant neuromodulation strategies involving NHPs and humans.

## Acknowledgements

The authors would like to thank Rafaella Mínea Riszt for her valuable technical contribution during equipment preparation and data acquisition.

The authors would like to express their gratitude to ADOTT Solutions Ltd., especially to Mr. Tamás Nagy for his vital contribution in the design and manufacturing the head-, and arm-fixation apparatus.

## Author contributions

A.P.: Conceptualization, Methodology, Investigation, Data Curation, Formal analysis, Visualization, Writing – Original Draft, Writing – Review and Editing;

B.K.: Conceptualization, Methodology, Software, Data Curation, Formal analysis, Visualization, Writing – Original Draft, Writing – Review and Editing;

E. K: Conceptualization, Methodology;

B.L.: Resources, Project administration, Funding acquisition, Writing – Review and Editing;

I.H.: Conceptualization, Methodology, Resources, Writing – Original Draft, Writing – Review and Editing, Supervision, Project administration, Funding acquisition.

